# On characterizing membrane protein clusters with *model-free* spatial correlation approaches

**DOI:** 10.1101/030718

**Authors:** Arun Shivanandan, Jayakrishnan Unnikrishnan, Aleksandra Radenovic

## Abstract

Spatial aggregation or clustering of membrane proteins could be important for their functionality, e.g., in signaling, and nanoscale imaging can be used to study its origins, structure and function. Such studies require accurate characterization of clusters, both for absolute quantification and hypothesis testing. A set of *model-free* quantification approaches — *free* of specific cluster models— have been proposed for this purpose. They include the radius of maximal aggregation *r_a_* obtained from the maxima of the empirical Besag *L*(*r*) – *r* function as an estimator of cluster size, and the estimation of various cluster parameters based on an exponential approximation for the Pair Correlation Function(PCF). However, the parameter identifiability and bias and scaling due to their *model-free* nature are not clear. In practice, the clusters might exhibit specific patterns, and the behavior of these estimators in such cases must be studied. Here, we theoretically analyze these approaches for a set of cluster models, and obtain information about their identifiability and bias. We find that the *ratio* between *r_a_* and true cluster size depends on both the true size as well as the number of clusters per unit area, or other corresponding parameters, in a model-dependent manner. In particular, *r_a_* scales with respect to the true size by a factor that can be arbitrarily large, depending on models and parameter values. For the method based on PCF approximation, for most models we analyzed, the ratios between approximate and true model parameters were found to be constants that depend only on models and independent of other parameters. For the models analyzed, this ratio was within ±100%. Our theoretical approach was validated by means of simulations. We also discuss some general issues in inference using second-order spatial properties. While precision could also be key, such information on identifiability and accuracy provides clarity on estimation, can lead to better inference, and can also fuel more accurate method development.

## Introduction

In cell biology and elsewhere, spatial aggregation or clustering is an interesting phenomenon, possibly with a functional role — e.g., the behavior of membrane proteins to form sub-micrometer clusters could be important for their functionality, such as in signaling (1–4). The origins, structure and function of spatial heterogeneity in membrane proteins are only being studied. Spatial location information, available from fluorescence and electron microscopic imaging, and recently from sub-diffraction limited fluorescence imaging such as Single Molecule Localization Microscopy(SMLM) techniques like Photo-Activated Localization Microscopy (PALM) and STochastic Optical Reconstruction Microscopy(STORM)(5–7), are key to such studies (8–11). Accurate characterization of clustering — the strength, scale and density of clustering is an important part of these studies, whether for relative comparison between different systems, perturbation conditions and to test hypothesis (such as the relative importance of lipid rafts and actin cytoskeleton in membrane protein clustering (12), or the possible mechanisms of early T-cell signaling (8)), or even for absolute quantification (such as the size of clusters in a particular cell type in a particular condition, and the number of molecules in them).

A number of methods have been used to characterize the clusters from imaging data (13–15). While most of these were aimed at characterizing membrane protein clusters, many of them were used to characterize other systems in the cell that exhibit clustering. The methods can be broadly categorized into two: (1) clustering or segmentation to identify the clusters, followed by their characterization; and (2) spatial statistics approaches based on a second-order spatial summary statistic such as Besag *L*(*r*) — *r* function or the Pair Correlation Function *g*(*r*). These second-order functions can be used for comparison of clustering at different scales and between different experimental systems and perturbations, and estimators based on these functions can be used for ensemble cluster parameter estimation. In general, they have a few advantages over many of the segmentation approaches: they can detect interactions at multiple spatial scales, can work with both dense and sparse point patterns, often have direct physical interpretations (16), and are amenable to rigorous extensions incorporating error models, crucial in the case of nanoscale imaging (16-18). Also, in the case of SMLM, the notion of spatial point patterns align well with the nature of its point localization readout. In practice, a major convenience of using such methods have been that they estimate ensemble functions at different scales and the various cluster parameters for a whole dataset, making comparative studies easy in systems where variability within cluster sizes are not important.

The simplicity in parameter estimation is in no small part aided by the *model-free* nature of some of these approaches. Two spatial statistics based estimators of cluster parameters based on these functions widely reported in the nanoimaging and protein cluster analysis literature, 1) the radius of maximal aggregation *r_a_* (13, 15, 18–27), the radius value corresponding to the maxima of the empirical *L*(*r*) — *r* function, as an estimator of cluster size(length scale); and 2) the *model-free* functional approximation of *g*(*r*) as an exponential function (9, 16, 28–32), leading to estimators of cluster size, amplitude or strength and number of molecules per cluster, are not concerned with the underlying spatial distributions, such as the shapes of clusters and the distribution of molecules in them. Effects due to differences in underlying spatial distribution are either ignored or approximated, effectively making the estimation process free of underlying cluster processes, or *model-free.* The *model-free* nature of these functions vary — *r_a_* does not contain any model of clustering whereas the PCF approximation is a generic function independent of specific models.

However, clusters observed through bio-imaging could be of different shapes, depending on the underlying physical mechanism. In the case of SMLM imaging, e.g., the clusters formed due to photoblinking are reported to have a Gaussian (9) or Cauchy peak shape (32), depending on the photon count distribution within the cluster. It is plausible to model internalization in circular or spherical bodies with a hard-core process (a disk in 2D). Analysis methods often assume Gaussian shapes for membrane clusters (14, 18). (33, 34) have suggested modeling membrane protein distributions using 2D-Ising model, to account for phase transitions and criticality. It is not clear how the parameter estimation approaches that are *model-free* are biased or scaled due to these different underlying true cluster processes. Also, such *model-free* approximations also raise the question of identifiability: e.g., can the size (i.e. length scale) parameter of model-free approaches be mapped exclusively to the size parameter of the true process, independent of other parameters, such as number of clusters per unit area or cluster density or amplitude? If the estimated size parameter is dependent on both the size and amplitude parameters of the underlying true process, one must account for it during the comparative analysis of cluster sizes, as it may not accurately reflect the true differences in size, being affected by amplitudes as well. Other point pattern based parametric methods (24, 35) also have to deal with similar issues. The influence of shape and geometry in estimation is observed in other fluorescence based technologies as well (36).

Some clues have been obtained from simulation studies. Kiskowski et al (19) studied the relation between the true radius of disk clusters *R* and estimates of *r_a_* by means of simulations, and derived important insights — such as *R* ≤ *r_a_* ≤ 2*R*, and a qualitative dependency of *r_a_* on separation between clusters. However, since the study was based on simulations, with a limited set of parameters and models (only disk clusters), the understanding is limited, and the possibilities of generalization are not clear. Lagache et al (24) performed a theoretical analysis of a similar estimator — maxima of the *K*-function normalized with its variance — for disk shaped clusters, and reported a simpler, constant relation *r_a_/R* = 1.3. Such a relationship would have been convenient, however its generality in terms of models and parameters is not clear. No studies of the bias introduced by the approximate model of *g*(*r*) has yet been reported, to the best of our knowledge.

Note that the accuracy or bias of an estimator cannot be improved by repeated measurements, unlike its precision. By definition, bias affects absolute quantification, e.g. an estimator of single molecule counts that is biased by 50% lower than the true counts affects absolute quantification. The same is the case regarding their use as relative comparisons: the need to account for biases might become important if (1) the parameters are not separately identifiable or (2) involves scaling that are model dependent and the comparisons involve different models.

In this work, we explore, with theoretical rigor, the bias in parameter estimation and the questions of identifiability introduced by these *model-free* approaches. We consider a number of spatial cluster processes whose theoretical *g*(*r*) and *L*(*r*) – *r* are known, and then derive the relation between the parameters of the *model-free* approaches (such as *r_a_*) and the true process parameters (e.g., the cluster size parameter *r_t_*). We find that, in general, for a large class of clustered point patterns, the ratio *p* of the radius of maximal aggregation *r_a_* and the size parameter of the true process *r_t_*(*p* = *r_a_/r_t_*) can be derived as an implicit function of two cluster parameters: *r_t_* and the number of clusters per unit area (*κ*). We also find that it possible to derive a theoretical lower bound for *p*, given a cluster model following some basic assumptions. We validate the theoretical results with simulations. We also perform similar analysis for the statistic presented in (24), to report a more complex relationship between the true cluster size and the estimator. Then, we investigate the bias due to the exponential approximation model of *g*(*r*), for all the models listed. By minimizing the Least Square Distance between the true and approximate PCFs, we obtain scaling laws between the approximate model and the true model parameters, and validate the approach by simulations. The extension for other cluster models are straightforward.

## Materials and Methods

### Background definitions

For a spatial point pattern in 2D-space, Ripley’s *K*-function is defined (37-39) as

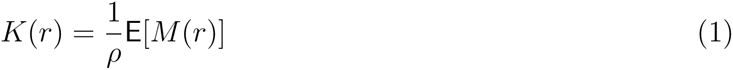

where *ρ* is the spatial density (average number of points per unit area), and *M*(*r*) is the number of other events within distance *r* of a randomly chosen event. The Besag *L*(*r*) — *r*, a measure of cluster strength at *r*, is then given by

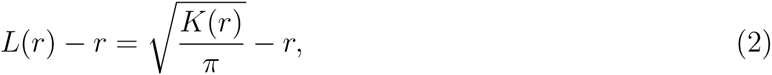

and the Pair Correlation Function by

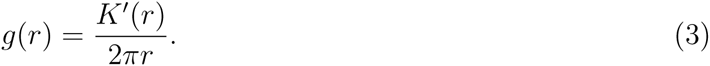

Alternative but equivalent definitions of PCF starting with the notion of spatial autocorrelation are also possible (9).

The radius of maximal aggregation, *r_a_* = arg max *L*(*r*) — *r*.

The function *g_a_*(*r*) = 1 + *a* exp(—*r/d*) has been proposed as a functional approximation for the Pair Correlation Function (PCF) of “2D-system of clusters with no predefined shape” (9, 28). The parameter *a* is the amplitude, a measure of point density in the clusters, and *d*, the correlation length, gives the radius of the cluster (9). For the PCF *g*(*r*), the average number of points per cluster can then be obtained as

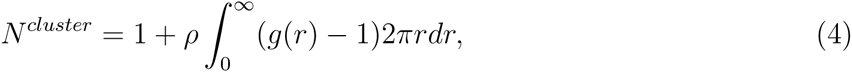

which is equal to *N_a_* = 2*πad*^2^*ρ* in the case of *g_a_*(*r*), where *ρ* is the average density of points in the area of analysis(9).

Theoretical expressions for *g(r)* and L(r) — r

In order to derive the theoretical expressions for *L*(*r*) — *r* and *g*(*r*) for different cluster models, it is useful to focus on a class of spatial cluster processes, known as Poisson cluster processes, or Neyman-Scott processes(details in (38, 39)), which are generated in the following way. First, a set of *parent* points are created, following a spatial Poisson process (complete spatial randomness) with density (intensity) *κ*. Then, *S* number of points are distributed around each *parent* point according to the i.i.d bivariate PDF *f_pdf_* (.), *S* following some i.i.d distribution with mean *μ*. These *offspring* points form the clustered point pattern. Such simple spatial cluster models that consider different shapes of clusters provide a starting point for the theoretical analysis of estimators. The analysis of Ising model in the later sections provide a more physical example.

Assuming *f_pdf_* (.) to be radially symmetric, let the PDF of the distance *r* between two offspring points within a cluster is given by *h_d_*(*r*) and its Cumulative Distribution Function (CDF) by *H_d_*(*r*). Then(38):

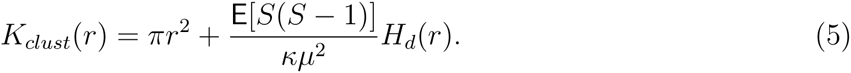

The density of the point pattern will be *μκ*. When *S* ~ *Poiss*(*μ*), since *E*[*S*(*S* — 1)] = *μ*^2^ for Poisson distribution, (5) reduces to

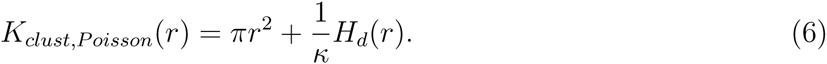

The derivation in case of other distributions for points per cluster is straightforward. In the case of geometric or exponential distribution of *S*, behavior often observed in nanoimaging (40-42), 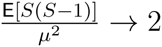 for *μ* ≫ 1.

Note that *H_d_,* being the CDF, is monotonic and non-decreasing. The corresponding PCF 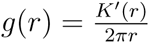 becomes:

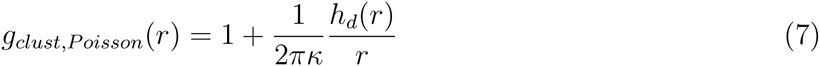

The PCF and *K*-function for different cluster shapes are given in Table 1, and the shapes of their PDFs are given in Supporting Material. Note that disk clusters contain points distributed uniformly at random within a circle (disk), a process known as Matérn cluster process in spatial statistics (the case of Gaussian cluster shapes is known as Thomas process). Also note that *r_t_* is defined differently for different cluster models: for a disk cluster, *r_t_* = *R*, the true cluster radius, whereas for Gaussian clusters, we set *r_t_* = *σ*, the true standard deviation (the full list can be found in Table 1). We also add the physical Ising model to the compilation, since it is one of the models that has been proposed for membrane protein clustering (16), even though it is not a Neyman-Scott process. Also, note that the exponential approximation *g_a_*(*r*) has the same shape as the variance Gamma function model(varGamma) in Table 1, pointing at the non-uniqueness of *g*(*r*) shapes and the difficulty of identifying cluster models from data based on their PCF shapes.

**Table 1.** Cluster models used for analysis.^†^. 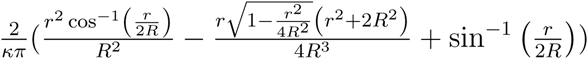. Also, for disk model, the functions provided here are for *r* ≤ 2*R*, for *r* > 2*R*, it is 0. Note that for disk, *g*(*r*) = 1 at *r* ≥ 2*R*, which provides a simple estimator for *R*.

| Model ($r_t$) | $g(r) - 1$ | $K(r) - \pi r^2$ |
| --- | --- | --- |
| Gaussian ($\sigma$) (39) | $\frac{1}{4\pi\kappa\sigma^2} \exp(\frac{-r^2}{4\sigma^2})$ | $\frac{1}{\kappa}(1 - \exp(\frac{-r^2}{4\sigma^2}))$ |
| disk ($R$) (39) | $\frac{2}{\pi^2 R^2 \kappa} (\cos^{-1}(\frac{r}{2R}) - \frac{r}{2R} \sqrt{1 - \frac{r^2}{4R^2}})$ | † |
| Cauchy ($\omega$) (45) | $\frac{1}{8\pi\omega^2\kappa} (1 + \frac{r^2}{4\omega^2})^{-3/2}$ | $\frac{1}{\kappa}(1 - \frac{1}{\sqrt{1 + \frac{r^2}{4\omega^2}}})$ |
| variance Gamma<br>$\nu = 1/2$ ($\eta$) (46) | $\frac{1}{2\pi\eta^2\kappa} \exp(-r/\eta)$ | $\frac{1}{\kappa} \left(1 - e^{-\frac{r}{\eta}} \left(1 + \frac{r}{\eta}\right)\right)$ |
| Ising (16) | $a_I r^{-1/4} \exp(-r/\xi)$ | $2\pi a_I \xi^{7/4} \left(\Gamma\left(\frac{7}{4}\right) - \Gamma\left(\frac{7}{4}, \frac{r}{\xi}\right)\right)$ |

### Effect of background

To model a monomer fraction or background, a spatial Poisson distributed monomer point pattern can be superimposed to a purely clustered process, such that the purely clustered fraction of points is *β*. The resulting *K*-function and PCF can be obtained using the expression for superposition of two independent point processes (38). In the case of a clustered process with *g*(*r*) = 1 + *Bv*(*r*), superposition with such a background process results in the PCF:

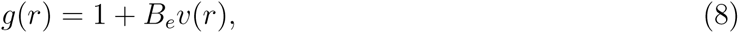

where *B_e_* = *Bβ*^2^, *β* being the purely clustered fraction (38). Expressions for *K*(*r*) and *L*(*r*) – *r* undergo similar scaling in parameter. It can be noted that the shape of the function remains the same as the purely clustered process, the change in parameter *B* being the only change, again pointing at the non-uniqueness of PCF shapes, and the quadratic effect of background on the function(note the effect on (4)).

### Simulation and analysis details

All simulations were done in `R`, using the `spatstat` library (43). Simulations of cluster processes were done with standard library functions, such as `rThomas` and `rMatClust`. Parameter estimation by minimum contrast method was done using `kppm` function, and using parameters “Thomas” and “VarGamma”. Analytical derivations were performed with the help of symbolic algebra software [Mathematica(Wolfram Research, USA)].

## Results

### Estimation based on radius of maximal aggregation

*Theoretical expressions for radius of maximal aggregation* Here we analyze the relationship between the radius of maximal aggregation, defined as *r_a_* = argmax *L*(*r*) – *r*, as a function of true cluster parameters, for the class of clustered point patterns with *K*-functions of the form

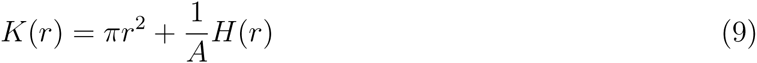

where *h*(*r*) = *H*′(*r*) and *A >* 0, such as the ones introduced in the Methods section and Table 1. Then, *L*′(*r_a_*) – 1 = 0, *L*′(*r_a_*) = 1 = ⇒ *K*′(*r_a_*)^2^ = 4π*K*(*r_a_*), using (2). Substituting in (9), we obtain

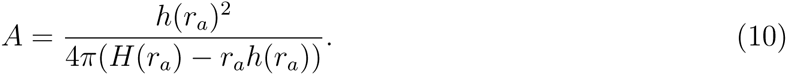

That is, *r_a_* depends on *A* in general, as *A* is not a parameter of *H* and *h*. (10) can be used to obtain a relation between 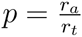 for all the models listed in Table 1, where *r_t_* is the cluster size parameter of the true process. The results are given in Table 2(more details in Supporting Material). It is possible to write the relationship 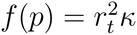 for all the Poisson cluster processes discussed. In the case of the Ising process, the corresponding relationship is of the form 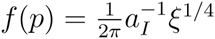. The derivation of p in the case of a power-law PCF is given in Supporting Material.

**Table 2.** Exact expressions for the radius of maximal aggregation *r_a_* for different cluster models.

| Cluster model | Expression for $p = r_a/r_t$ | Theoretical lower bound for $p$ (to 5 digits) |
| --- | --- | --- |
| Gaussian ($p = r_a/\sigma$) | $\kappa\sigma^2 = \frac{e^{-\frac{p^2}{4}} p^2}{8\pi(-p^2 + 2e^{\frac{p^2}{4}} - 2)}$ | 2.24181 |
| Disk ($p = r_a/R$) | $\kappa R^2 = \frac{p^2 \left( p \sqrt{4 - p^2} - 4 \arccos(\frac{p}{2}) \right)^2}{\pi^2 \left( \sqrt{4 - p^2} (3p^2 - 2) p - 8p^2 \arccos(\frac{p}{2}) + 8 \arcsin(\frac{p}{2}) \right)}$ | 1.29564 |
| Cauchy ($p = r_a/\omega$) | $\kappa\omega^2 = \frac{p^2}{\pi(p^2 + 4)^{3/2} ((p^2 + 4)^{3/2} - 4p^2 - 8)}$ | 2.54404 |
| varGamma ($p = r_a/\eta$) | $\kappa\eta^2 = \frac{p^2}{4\pi(\exp(2p) - \exp(p)(p^2 + p + 1))}$ | 1.79328 |
| Ising ($p = r_a/\xi$) | $\frac{1}{2\pi} a_I^{-1} \xi^{1/4} = \frac{\exp(-2p)p^{3/2}}{4\pi(-\exp(-p)p^{7/4} - \Gamma(\frac{7}{4}, p) + \Gamma(\frac{7}{4}))}$ | 1.37220 |

Note that the expression for *p* (and hence *r_a_*) is independent of the number of points per cluster (*μ*) if the expressions for *K*-functions are independent of it. Figure 1 shows the contour plot of *p* vs *κr_t_*. Thus, we can establish that theoretically, the ratio between the radius of maximal aggregation and the true size parameter is dependent on both the true size parameter as well as the number of clusters per unit area. In fact, the singularity at *H*(*r_a_*) − *r_a_h*(*r_a_*) = 0 provides a minimum bound for *p* for all the models analyzed, and is also shown in Table 2(see Appendix for a proof). The lower bound so obtained is a fundamental characteristic of the cluster model’s theoretical *L*(*r*) − *r* functions. The existence of a lower bound for *r_a_* for any cluster model with *K*-function of the form in (9) can be proved theoretically given some basic assumptions on *h*(*r*)(see Appendix).

**Figure 1.**
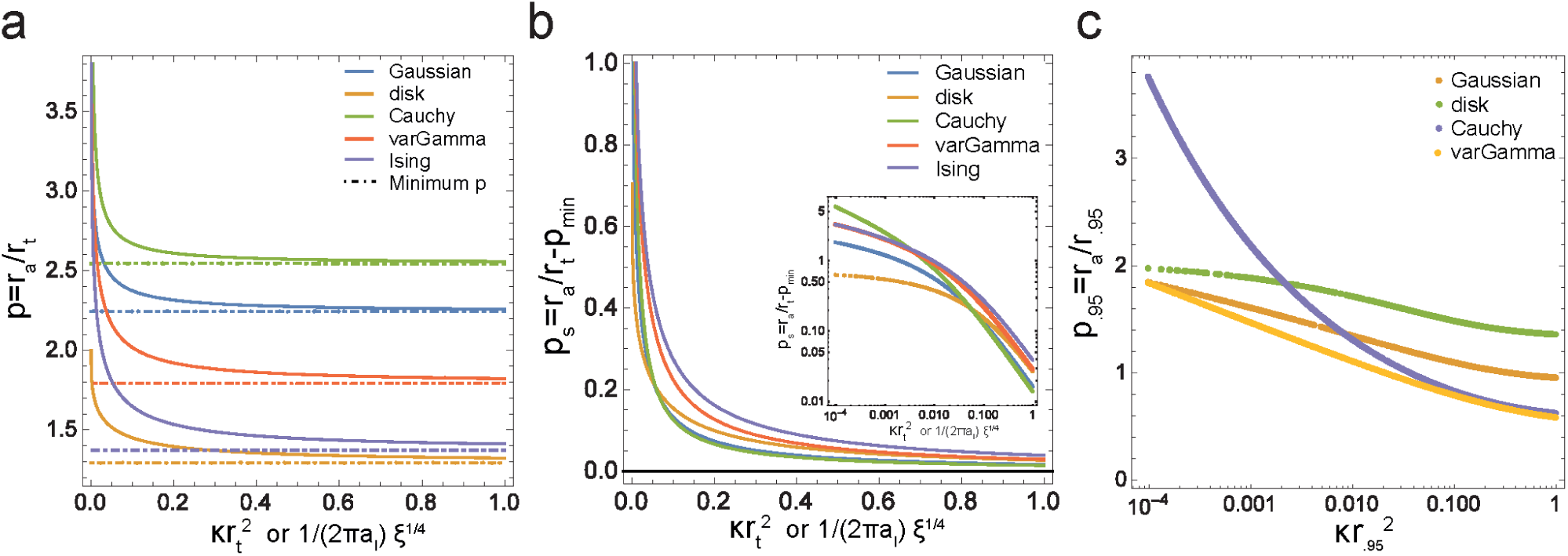
Relation between the radius of maximal aggregation and true cluster size. a For different cluster models, the relation between the ratio of radius of maximal aggregation *r_a_* and cluster size parameter of the true process *r_t_,* as a function of the number of clusters per unit area *κ* and *r_t_*. The minimum *p* value is obtained by exploiting the singularity in (10), given in Table 2 b Plots in a after translating by the minimum *p* and in log-log scale (inset). Note the partial power law like shape. **c** *p*._95_, the ratio between *r_a_* and *r*._95_, the latter being the true scale within which 95% of all clustered points lie, plotted against *κr*^2^_95_. It can be seen that the relationships are model dependent. Note that for a sample with 10 clusters per *μm*^2^ and *r*. _95_ = 20*nm, κr*^2^_95_ = .004.

The *p* = *r_a_/r_t_* for different processes cannot be directly compared, as the size parameter of the true process *r_t_* is defined differently for them. Now, it can be seen that the lower bound for *p* is model dependent. For the disk model, e.g., 1.29564 < *p* < 2, whereas, for Gaussian model, 2.24181 < *p* < ∞. A more comparable measure would be *r_q_*, the (true) scale at which *q* fraction of the points are expected to lie for a particular distribution, typically obtainable in the form *r_q_* = *u_q_r_t_*, such as the case of *r*._95_ = 2*σ* in the case of 1D Gaussian distribution. This would correspond to the ratio *p_q_* = *r_a_*/*r_q_* = *p*/*u_q_*, and 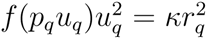. Considering the case *q* = 95%, the values for *u*._95_ and the lower bounds for *p*._95_ for different distributions is given in Supporting Material, and the plot *p*._95_ *vs κr*^2^_95_ is shown in Figure 1c. It can be seen that *p*._95_ is dependent on both the model as well as both the number of clusters per unit area and the true cluster size.

The systematic relationship established between *p* (or *p*._95_), *A* and *r_t_*, clarifies the bias and identifiability issues in estimation. The results agree with (19), and provides a tighter (theoretical) lower bound for disk clusters. The approach can also explain the qualitative influence of inter-cluster distance on *r_a_*, observed by (19), through the dependency of *p* on *κ*, along with the relative influence of *κ* and *r_t_*. The dependency of *p* on other cluster parameters and the cluster model means that the estimator could be a poor choice as a comparison tool between different experiments, if there is a possibility that the cluster model, *κ* or *r_t_* are different — unless there is a large enough difference in estimated *r_a_* for different experiments.

*Validation with simulations* To establish the validity of the theoretical derivation obtained in previous section (shown in Table 2) we performed a Monte Carlo simulation study. In addition to information about the accuracy of radius of maximal aggregation(the subject of the theoretical study), it also provides information about its precision as an estimator.

Clustered point patterns, belonging to either Gaussian or disk clusters, were simulated in a unit square, for varying *κ* and *r_t_*. The theoretical value of *p* for a given *κ* and *r_t_* were obtained by solving the analytical expressions in Table 2, and was compared to 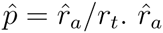 was obtained from the empirical maximum of the *L*(*r*) − *r* curves. The results are shown in Figure 2(also see Supporting Material Figure 1). The mean value of 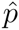 from simulations broadly agree with the theoretical results, though the deviation increases with increasing 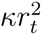(see also the Mean Squared Error in Figure 2c,d). This is probably the result of increasing number of clusters per unit area (increasing *κ*) or having larger clusters within the unit square used in the simulations (increasing *r_t_*), both resulting in overlapping clusters, resulting in deviations from theoretical framework based on a particular cluster model. In fact, it can be seen that the deviation is most influenced by increasing radius(Figure 2c,d).

**Figure 2.**
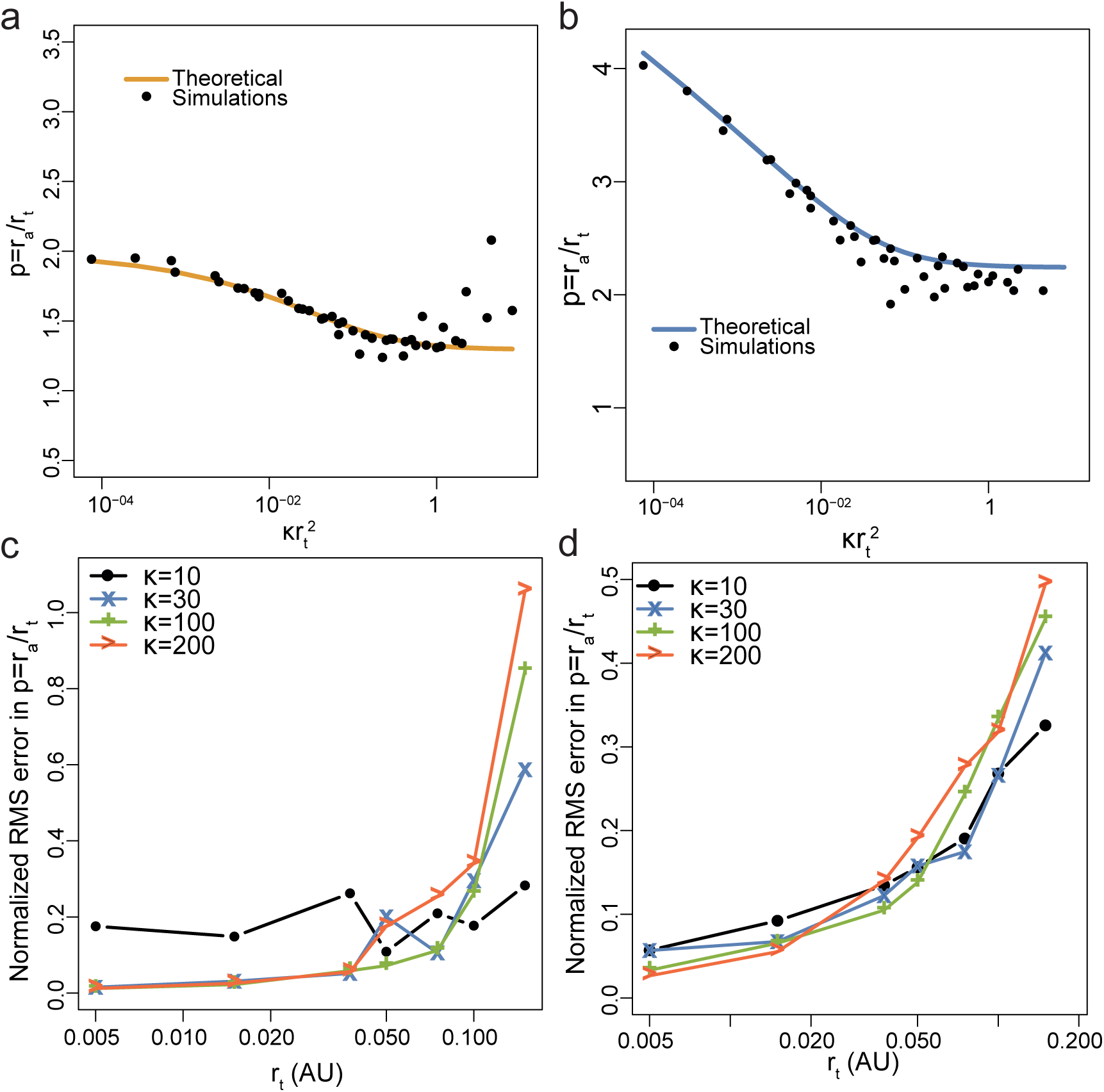
Comparison of theoretical results on *p* = *r_a_*/*r_t_* with that from simulations **a,b** Results from theory (solid curve) as well as simulations on unit square window(dots), for disk and Gaussian clusters respectively. Only the mean value from 100 simulations are shown, for clarity, and the plot with error bars can be seen in Supporting Material Figure 1. It can be seen that in both disk and Gaussian cases, the mean values from simulations deviate from the theoretical values with increasing 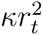. **c,d** The Root Mean Squared error, normalized by the theoretical value, for disk and Gaussian clusters respectively, plotted against *r_t_*. The colors denote different *κ* values. It can be seen that the error values are highly influenced by *r_t_*.

*Case of normalized K-function* In (24), the normalized statistic 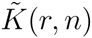 was proposed, given by 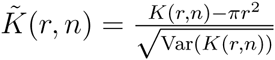, where

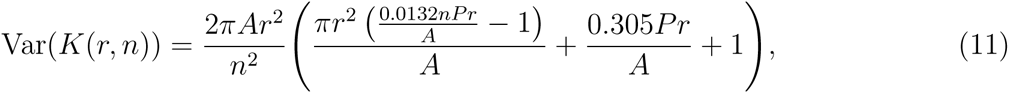

where *A* is the area and *P* the perimeter of the observation window, and *n* the number of points.

For disk process, they use, similar to the expression in Table 1:

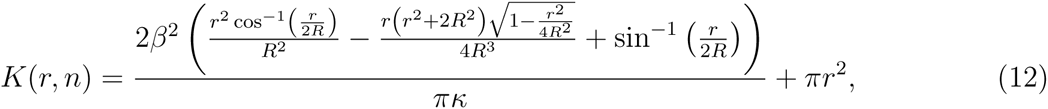

where *κ* is the number of clusters per unit area, and *β* the clustered fraction.

The radius of maximal aggregation 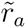 for 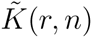 is then obtained by setting 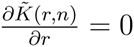. Using numerical approaches, they obtained the constant relation 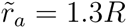.

In our hands, 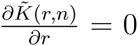 for a square observation window(for simplicity) resulted in a more nuanced situation, as shown in Figure 3(details in Supporting Material). We found that 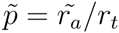 depends on the number of points *n* and the ratio *m* between the side length of the square observation window and the true size parameter *r_t_*, and converges to a maximum value at large *m*, which is approximately equal to the minimum values obtained in the case of *r_a_* based on *L*(*r*) − *r*. For example, in the case of clusters with *R* = 20*nm* with an area of analysis of size 10*μm*, then *m* = 500, and 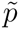 is close to the maximum value (Figure 3), and hence a constant(1.296 in the case of disk clusters, approximately equal to the factor of 1.3 obtained in (24)). On the other hand, if the area of analysis was smaller, say 1*μm*, then *m* = 50, and 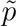 depends critically on *n*(Figure 3). The dependency of 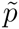 on *n*, in contrast with *p* in the case of *L*(*r*) − *r*, is because 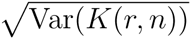 is non-linearly dependent on *n*, whereas the expression for *K*(*r*, *n*) (and *L*(*r*) − *r*) is independent of *n*. Note that 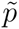 is independent of *κ* and *β*, unlike the case of *r_a_* and *L*(*r*) − *r* presented in the previous section.

**Figure 3.**
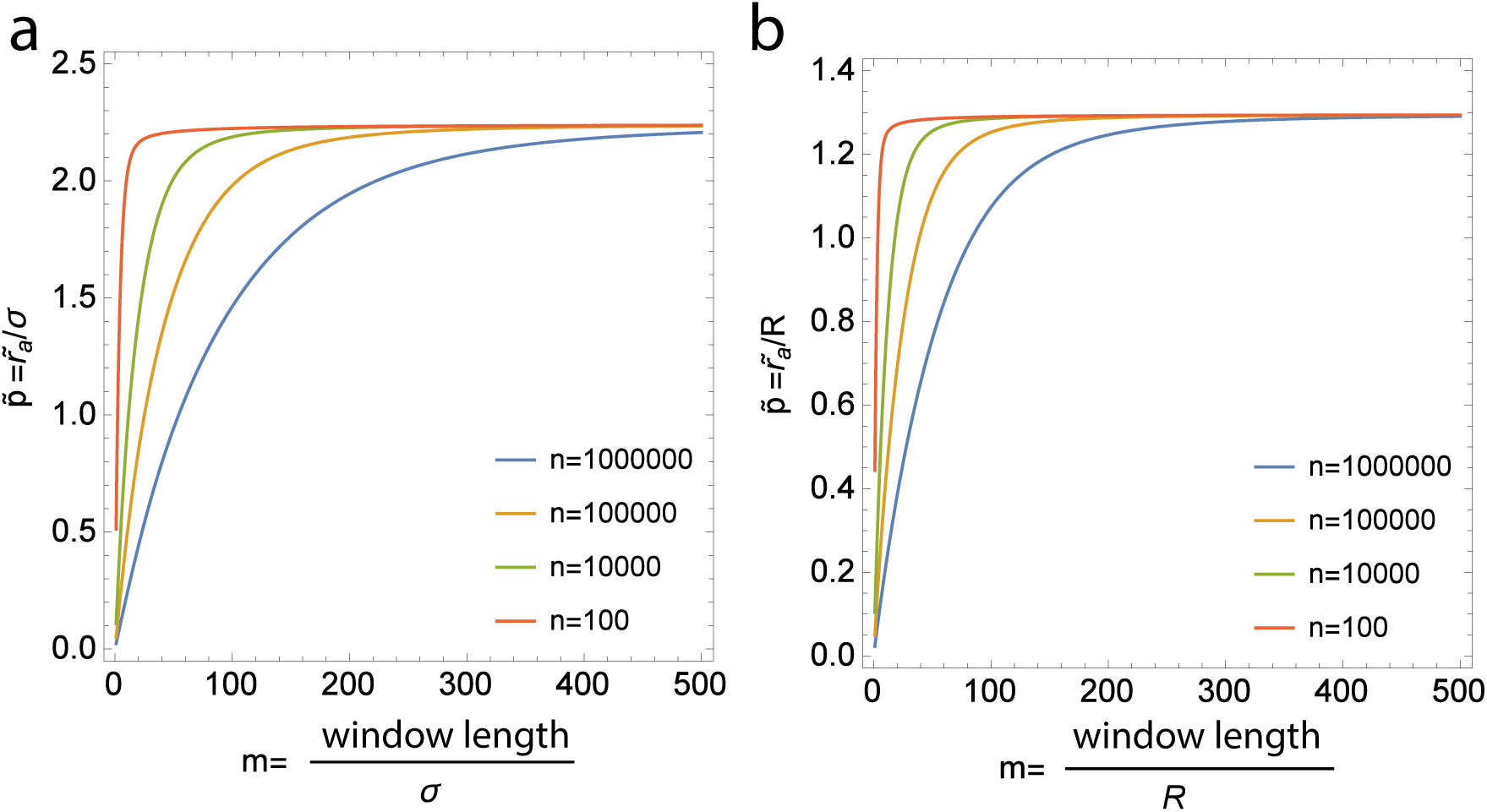
Results in the case of normalized *K*-function 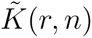. **a** Gaussian clusters, **b** disk clusters. In the case of 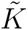, the 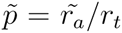 depends on the number of points *n* and the ratio *m* between the size of the observation window (side length of a square in this case) and the true size parameter *r_t_*, and converges to a maximum value at large *m*, which is approximately equal to the minimum values obtained in the case of *r_a_* based on *L*(*r*) − *r.* Note that in the case of clusters with *R* = 20*nm* with an area of analysis of size 10*μm*, *m* = 500, and 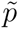 is close to the maximum value, and hence a constant. On the other hand, if the area of analysis was smaller, say 1*μm*, *m* = 50, and 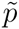 depends on *n*.

### Estimation based on Pair Correlation Function

We now consider another estimator that has been suggested for estimating cluster parameters, the approach based on fitting Pair Correlation Functions. As discussed in the section Methods, the theoretical PCF is not unique for a cluster model, and its signature shape and sensitivity are often not sufficient to identify the models(Figure 4), not the least because each experiment provides a realization of a stochastic process, with the observed statistic approaching the theoretical one only as *n* → ∞. Model selection based on Monte Carlo (MC) rank tests (38, 39) — ranking the empirical statistic value among the values of the statistic from MC simulations based on estimated parameters — based on PCF or the related *K* or *L*(*r*) − *r* functions is not sound, if the same function was used for parameter estimation(38). The standard method in this case is to perform MC rank tests with a statistic that is different from the one that was used for parameter estimation, e.g., the nearest neighbor distribution function if the PCF was used for estimation. However, the approach is known to have low statistical power (39), and we too had similar experience during preliminary attempts to identify the cluster models from simulations and SMLM data (results not shown). Therefore, the *model-free* functional approximations such as *g_a_*(*r*) = 1 + *a* exp(−*r*/*d*), proposed as part as the PC-PALM method, have much appeal.

**Figure 4.**
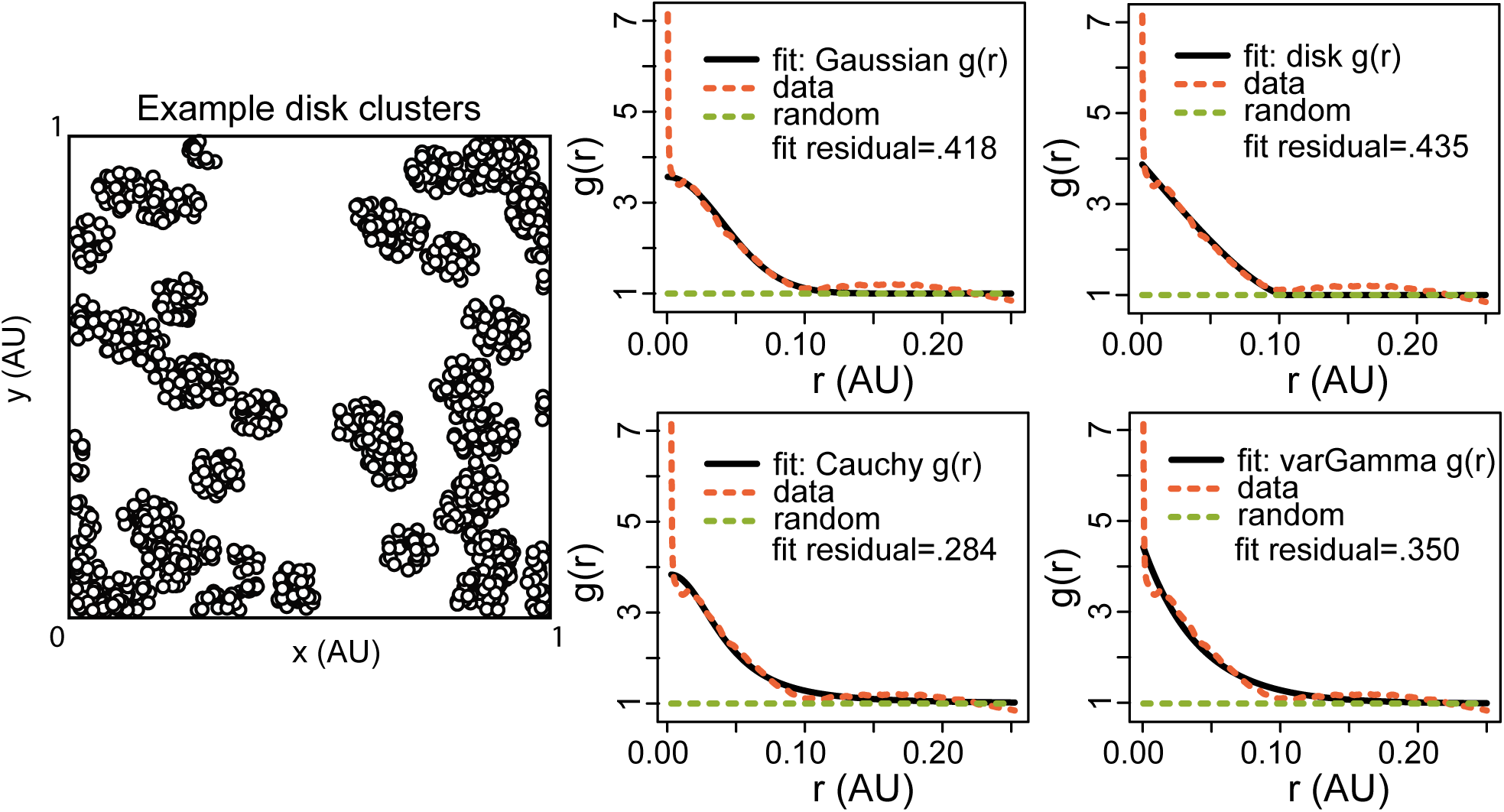
Demonstrative example of fitting model PCFs to the empirical PCF of a disk point pattern. The empirical PCF of the point pattern in the left is calculated, and is fit to the theoretical PCFs of various cluster processes. Fit results 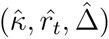, 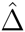 being the value of the objective function for the best fit parameters, called fit residual: Gaussian (38.11, .028, .418), disk (40.64, .052, .435), Cauchy (21.55, .051, .284), varGamma (27.86, .040, .350)), whereas the true values of the disk point pattern are (*κ* = 50, *r_t_* = *R* = .05). Note that 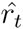 is defined differently for different processes (Table 1). The Cauchy distribution is found to have the best fitness, whereas the disk one — the true model — has the worst. The *p* = *r_a_*/*r_t_* corresponding to disk distribution, with the estimated parameters above is 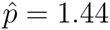. The maxima of *L*(*r*) − *r* is at 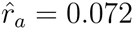, providing a 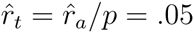, equal to the true *R*.

Here, we derive a measure of bias in parameters introduced by this approximation, given a true model. We aim to find the relations *m* = *d*/*r_t_*, *n* = *a*/*a_t_* and *l* = *N_a_*/*N_t_*, given a true model for the PCF in the form *f* (*r*) = 1 + *a_t_v*(*r*, *r_t_*). Here, *N_a_* and *N_t_* are the average number of points per cluster corresponding to the approximate model and the true model respectively, as per (4). Given a specific model for *f*(*r*), we find the relation between parameters in the case of the fit that provides the minimum (Least) Squared Error *E*, i.e.,

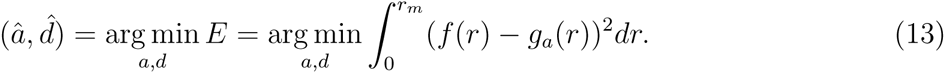

Note that the Least Squares criteria was used in original PC-PALM papers for parameter estimation (9, 44). If *E* has a minima at 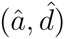, then 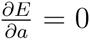 and 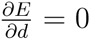 at 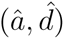, which can be solved to obtain expressions for 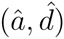. Measures of *m*, *n* and *l* can then be found using these.

We were able to obtain measures of *m*, *n* and *l* for all the cluster models described in Table 1, and the results are shown in Table 3 and the best fit PCFs can be seen in Figure 5a(details in Supporting Material). The *m*._95_ values: *m*._95_ = *d*/*r*._95_ = *m*/*u*._95_, *r*._95_ being the scale at which 95% of points are expected to lie, can also be obtained as constant scalar values, given by .63,.82,.38 and .28, for Gaussian, disk, Cauchy and varGamma models respectively.

**Figure 5.**
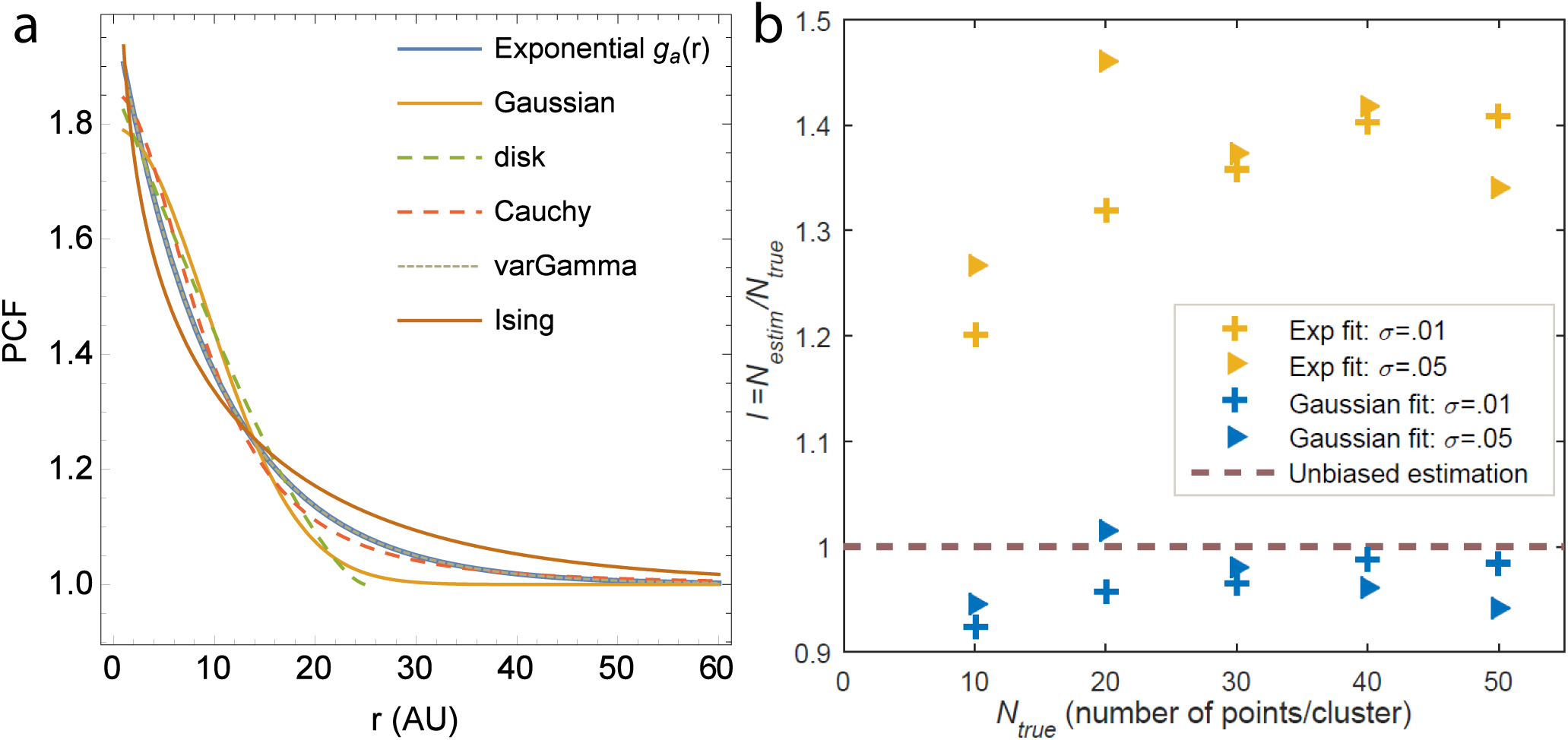
**a** Optimal Least Square Error fits for different models. For parameter values *a* = 1 and *d* = 10, the PCFs corresponding to different models in Table 1 is plotted, with the parameters scaled as per Table 3. For simplicity, only *r* ≥ 1 is shown. **b** Mean estimates of *N* (number of points per cluster) from fitting the empirical PCF of Gaussian clustered point patterns with (1) Gaussian PCF (2) the exponential approximation *g_a_*(*r*)(results from 20 simulations on a unit square window). The results broadly agree with the theoretical prediction of *l* = 1.48, approaching it with larger *N_true_.* A plot with error bars can be seen in Supporting Material Figure 5.

**Table 3.** Theoretical scaling for different cluster models, in using the exponential approximation for PCF and using Least Square Error criteria. *d*, *a*, *N_a_* correspond to the approximate PCF model *g_a_*(*r*) = 1 + *a* exp(−*r*/*d*). True parameters *r_t_*,*a_t_* and *N_t_* corresponding to the model PCFs of the form *f* (*r*) = *a_t_v*(*r*, *r_t_*) can be obtained from Table 1 and using 4. The minimum *r_m_* value, used in the calculation of the Squared Error *E* (in 13), for each model is as follows: Gaussian - 6*σ*, disk - 3*R* and Ising - 4*ξ*, and higher values for *r_m_* give the same results. In the case of Cauchy model *r_m_* = ∞ was used, and for varGamma any *r_m_* > 0 corresponds to the results in the table. The *m*._95_ values: *m*._95_ = *d/r*._95_, *r*._95_ being the scale at which 95% of points are expected to lie, are .63,.82,.38 and .28 respectively, for Gaussian, disk, Cauchy and varGamma models.

| Cluster model | $m = d/r_t$ | $n = a/a_t$ | $l = N_a/N_t$ |
| --- | --- | --- | --- |
| Gaussian | 1.54 | 1.26 | 1.48 |
| disk | .8 | 1.81 | 1.48 |
| Cauchy | 1.7 | 1.17 | .85 |
| varGamma | 1 | 1 | 1 |
| Ising | .5 | $2.15r_t^{-1/4} (\approx .38 - 1.44)$ | .59 |

For example, in the case of Gaussian shaped clusters, with the PCF given in Table 1, we obtain, for *r_m_* > 6*σ*, *m* = *d*/*σ* ≈ 1.54, *n* ≈ 1.26, *l* ≈ 1.48, with *m*._95_ = .63. The parameters can be either *upscaled* or *downscaled* — e.g., the number of molecules per cluster is overestimated by 50% by using *g_a_*(*r*) for estimation, whereas in the case of Ising process, it is underestimated by 40%. The overestimation/underestimation for all parameters is no more than by 100%, in all the models the approach was applied, except in the case of the amplitude parameter in the Ising model. In this case too, while the *a* parameter is dependent on both the true amplitude *a_I_* as well as true size parameter *ξ*, the effect is to the extend of *n* = .38 − 1.44 for *ξ* =5–1000nm, the case relevant in the case of membrane protein clusters.

For the models in Table 1, this means that (1) the scaling is either independent of other parameters or only mildly dependent (2) the theoretical scaling due to the exponential approximation is within 100%, in contrast with the radius of maximal aggregation, which can be several times higher (technically upto ∞) depending on models and parameter values.

We validated this theoretical approach by means of Monte Carlo simulations. We simulated Gaussian cluster processes in a unit square for different conditions, such as varying the numbers of points per cluster as well as cluster radius. The empirical PCF of these point patterns were fitted to both the theoretical PCF for Gaussian point patterns, as well as the functional approximation *g_a_* (*r*), and the various parameters estimated. The estimates for *N*, the number of points per cluster is shown in Figure 5b. It can be seen that the simulations agree with the theoretical prediction, with estimates using *g_a_*(*r*) being overestimated, whereas the fit to Gaussian PCF providing accurate results.

## Discussion

We have theoretically analyzed three spatial statistics based *model-free* methods for cluster parameters that have been proposed in the membrane protein imaging literature. They are: the radius of maximal aggregation based on Besag *L*(*r*) − *r* function and the radius of maximal aggregation based on normalized *K*-function, both primarily estimators for cluster size, and the estimation based on the functional approximation with an exponential function for the Pair Correlation Function, proposed in the PC-PALM method. We were able to derive the theoretical relation between the radius of maximal aggregation and the true cluster parameters, for a diverse set of models, along with a theoretical lower bound for it. Our results illustrate the fact that the ratio of the radius of maximal aggregation (in the *L*(*r*) − *r* case) to the true cluster size depends on the true cluster size as well as the number of clusters per unit area (or corresponding parameters, such as amplitude) for all the models considered. This dependence points to the difficulties of parameter identifiability using this technique, and also has implications in the interpretation of empirical *L*(*r*) − *r* curves. In the case of the Pair Correlation approach, we were able to derive the scaling laws between the parameters of the approximate model and the true model, based on the Least Square Error criteria. From both the identifiability point of view as well as the scale of bias (between the true process parameters and the estimators), it appears that the Pair Correlation approach performs better, at least for the models our approach was applied on. While only a limited set of models were analyzed here, they show the limits of the estimators, and extending the analysis to other models is straightforward.

Also, the analysis shows that it might be possible to obtain theoretical bounds for parameters given a set of candidate models, e.g. by taking the worst bounds among candidates, even though the specific candidate model for a system is not known or is difficult to be inferred. It also points to a possible approach to reducing the bias: by using non-parameteric models for the *model-free* PCF, although care must be taken against overfitting and also in interpreting the results. This work only deals with the accuracy limits of the estimators, their precision could also be important in practical applications, which must be analyzed separately. The results presented in this work are not limited to membrane protein clusters, and are applicable to any system with spatial clustering.

## Appendix

### Lemma 0.1.

*Let h :* 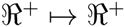 *be a unimodal differentiable function with a unique maximum at r_m_* > 0 *and a derivative satisfying h*′(*r*) > 0 *for* 0 ≤ *r* < *r_m_, and h*′(*r*) < 0 *for r > r_m_. Note: this is satisfied by all the models in Table 1.*

*Further assume that there exists r\** > 0 *that satisfies*

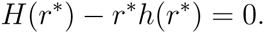

*Then the radius of maximal aggregation r_a_ ≥ r* where r_a_ is obtained as a solution to (10) for some A* > 0.

*Furthermore as A* → ∞*, we have r_a_* → *r*.*

#### Proof.

Define

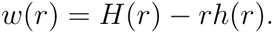

Clearly *w*(0) = 0 and the derivative satisfies *w*′(*r*) = −*rh*′(*r*).

From the properties of *h*′ we have *w*′(*r*) ≤ 0 for 0 ≤ *r* < *r_m_,* with strict inequality for 0 < *r* < *r_m_,* and *w*′(*r*) > 0 for *r* > *r_m_.* Hence

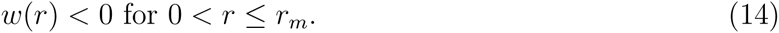

Since *w*(*r*\*) = 0 it follows that *r*^*^ > *r_m_.* Moreover since *w*′(*r*) is strictly positive for *r* ∈ (*r_m_*, *r*^*^], it follows that *w*(*r*) < 0 for *r* ∈ (*r_m_*, *r*^*^). Combining with (14) it follows that *w*(*r*) < 0 for *r* ∈ (0, *r*^*^).

Now, we know that *r_a_* satisfies (10) for some *A* > 0. Thus we must have *w*(*r_a_*) > 0 and hence it follows that *r_a_* ≥ *r*^*^.

Now consider the situation in which *A* → ∞. Define

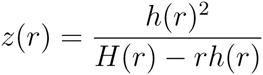

to denote the expression on the right hand side of (10) without the factor of 4*π* included. Since 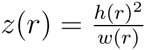 we know from the earlier analysis of *w* that *z*(*r*) ≤ 0 for *r* < *r*^*^ and *z*(*r*) ≥ 0 for *r* < *r*^*^. Now consider the derivative of *z*. We have

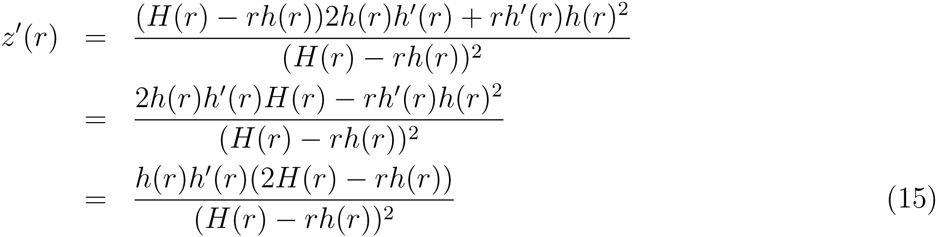

Now consider the function *q*(*r*) = 2*H*(*r*) − *rh*(*r*) for *r* ≥ *r*^*^. At *r* = *r*^*^ we have *q*(*r*^*^) = 2*H*(*r*^*^) − *r*^*^*h*(*r*^*^) = *H*(*r*^*^) > 0. Moreover the derivative of this function is *q*′(*r*) = *h*(*r*) − *rh*′(*r*) which is non-negative for *r* > *r*^*^ because *h*′(*r*) < 0. Thus *q*(*r*) > 0 for *r* > *r*^*^. This observation combined with the fact that *h*′(*r*) < 0 for *r* > *r*^*^ and (15) implies that *z*′(*r*) < 0 for *r* > *r*^*^. Thus we have that *z* is strictly decreasing in the interval (*r*^*^, ∞). Moreover *z*(*r*) → ∞ as *r* approaches *r*^*^ from above. Hence as *A* → ∞ the left hand side of (10) → ∞ and thus by virtue of (10) we must have *r_a_* → *r*^*^. □

## Author Contributions

A.S. conceived the project, performed the analysis, and wrote the manuscript, with inputs and guidance from J.U. and overall supervision from A.R. All authors read and approved the final manuscript.

## Acknowledgments

A.S. thanks Prof. Ivo Sbalzarini, Hendrik Deschout and Dileep Kalathil for early discussions and guidance. This work was financially supported by Swiss National Science Foundation grants No. 200021–125319 and No. 20021–132206. A.S. was funded by a PhD fellowship grant from NCCBI. The authors declare that there are no conflicts of interest.

## Supporting Material for *On characterizing membrane protein clusters with model-free spatial correlation approaches*

### 1 Derivation of 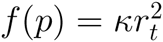 and similar expressions for *p*

Here we derive the relation in the case of Neyman-Scott process with Gaussian shaped clusters. The derivation in the case of other distributions are similar, starting from the expressions in Table 1, Main Text.

We start from the *K*-function for Gaussian shaped clusters:

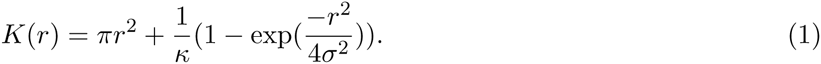

In the form 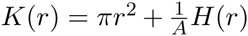 as in Main Text, this corresponds to 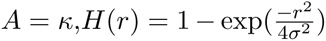 and 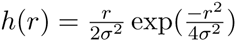. Substituting in the equation

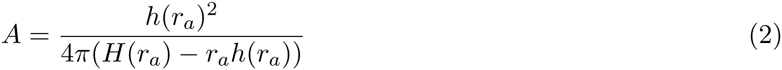

from Main Text and rearrangement will give the relation as in Table 2, Main Text.

### 2 95% scale for various models

These were found by solving the CDF 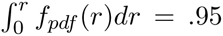 for *r*, where *f_pdf_* (*r*) is the radial probability density function for each model(1–3). In the case of Cauchy and varGamma models, marginal PDFs of *r* in polar coordinates were obtained from the bivariate PDFs in cartesian coordinates by standard transformation(multiplication by 2*πr*). The results are given in the following table, along with the 95% limits. *K_v_*(.) denotes the modified Bessel function of the second kind.

**Figure 1:**
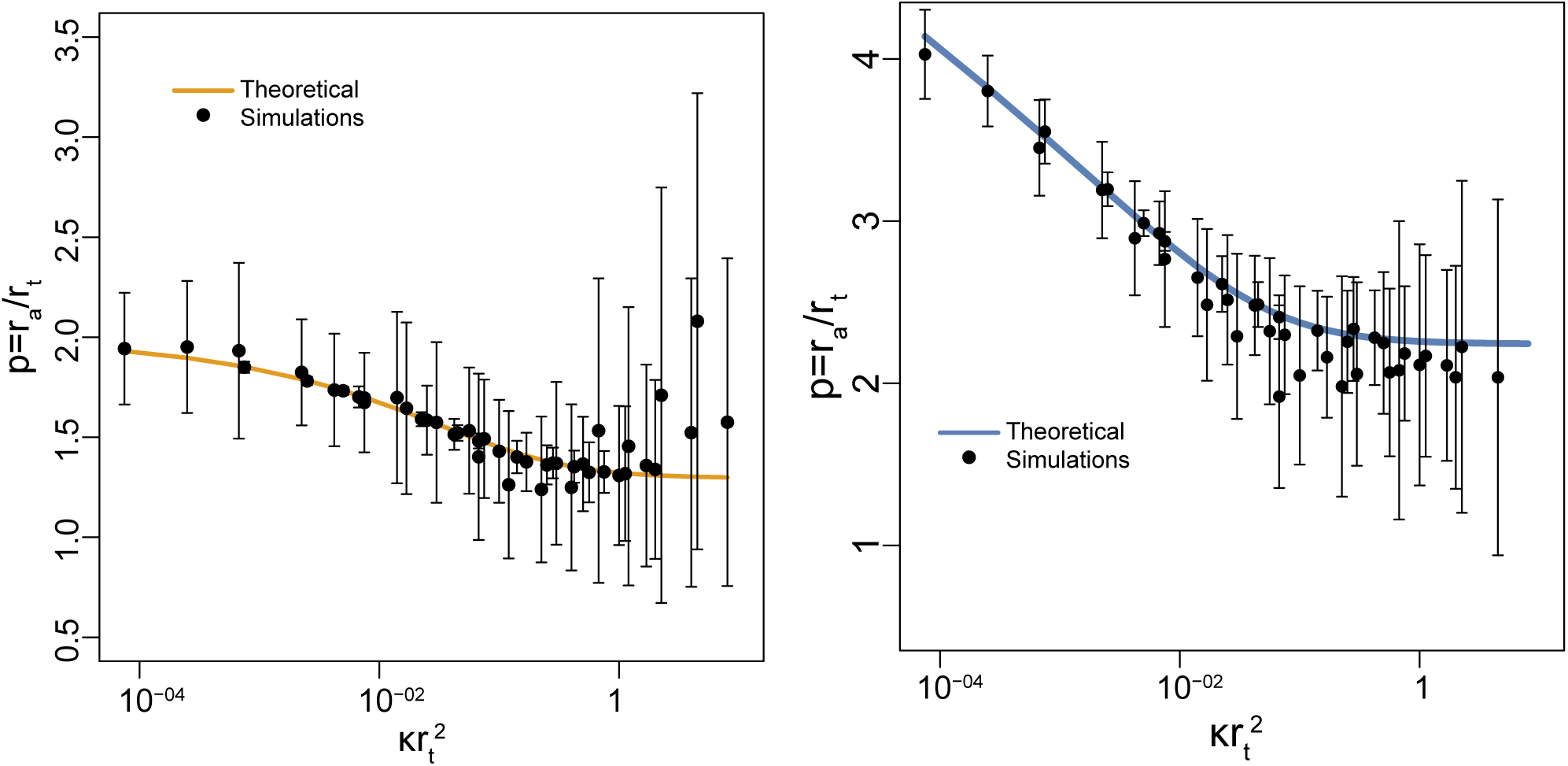
Comparison of *p* = *r_a_*/*r_t_* from theory and simulations. Figure 1 in Main Text with error bars(*σ*).

| Model | $f_{pdf}(r)$ | $r_{.95} = u_{.95}r_t$ | Lower bound for $p_{.95}$ |
| --- | --- | --- | --- |
| Gaussian | $\frac{r}{\sigma^2} \exp\left(-\frac{r^2}{2\sigma^2}\right)$ | $2.448\sigma$ | .914 |
| disk | $\frac{2r}{R^2}$ | $.975R$ | 1.329 |
| Cauchy | $\frac{r}{\omega^2} \left(1 + \frac{r^2}{\omega^2}\right)^{-3/2}$ | $4.469\omega$ | .568 |
| VarGamma | $\frac{\sqrt[4]{2}r^{3/4}K_{-\frac{1}{4}}\left(\frac{r}{\eta}\right)}{\eta^{7/4}\Gamma\left(\frac{3}{4}\right)}$ | $3.547\eta$ | .505 |

### 3 Radius of maximal aggregation in the case of 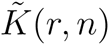 of Lagache et al

Setting 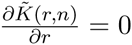 for disk clusters as discussed in Main Text, followed by routine manipulations lead us to the relation:

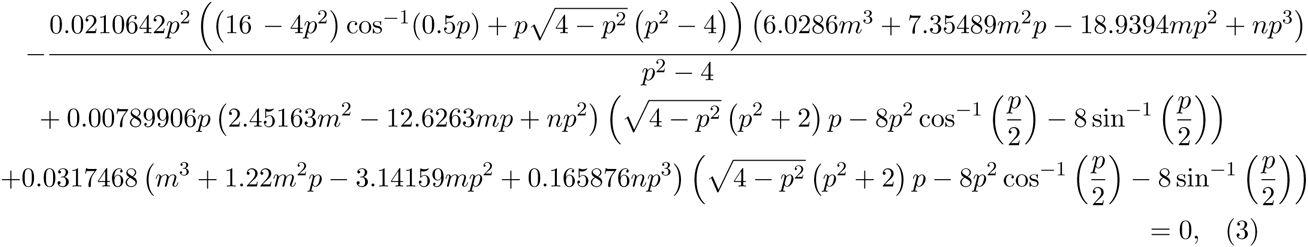

where 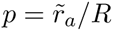, *m* = *side/R* where *A* = *side^2^*, *P* = 4.*side*.

The contour plot of *p* vs *m*, based on this expression, is shown in the Main Text, for different values of *n*.

In the case of Gaussian clusters, the relation is simpler:

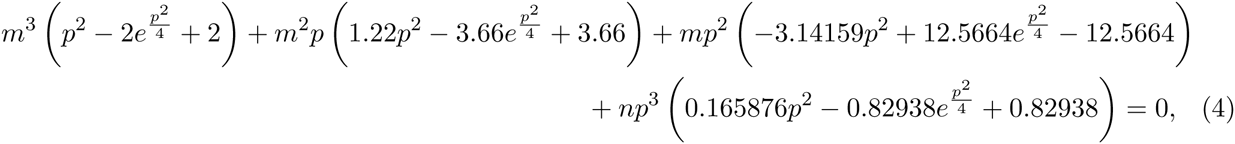

and the corresponding contour plot is provided in Main Text.

### 4 Derivation for bias in PCF based on Least Squared Error

We simply show the case for Ising model. Derivation for other models follow the same procedure. For *g_a_*(*r*) = 1 + *a* exp(−*r/d*) and *f* (*r*) = 1 + *Ar*^−1/4^ exp(−*r/D*), the Least Squared Error criteria gives:

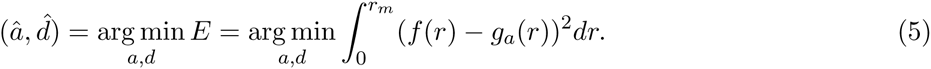

We obtain: 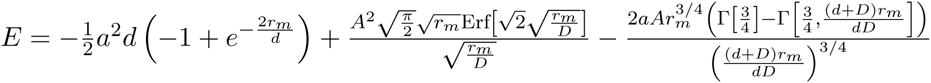

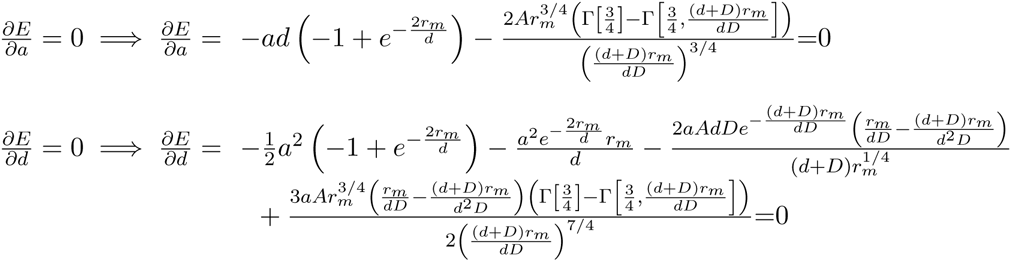

Solving both equations separately for *a* = *â*, we obtain:

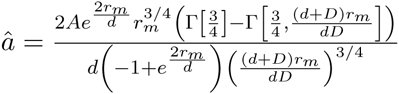

and,

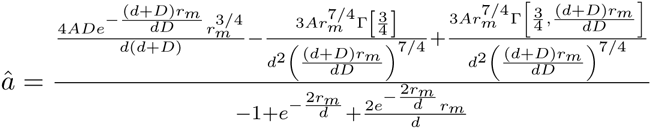

Equating both the above expressions of *â*, simplifying, and setting *m* = *d/D* and *k* = *r_m_/D*, we get:

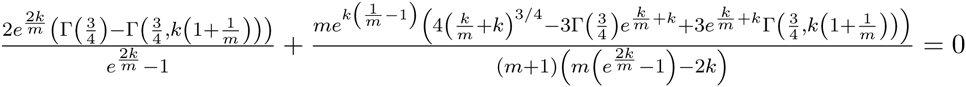

Note that this equation does not contain the amplitude parameters *a* and *A*. A contour plot of this equation is shown in Figure 2. For reasonably large values of *r_m_* (i.e., *r_m_ >* 2*D*), 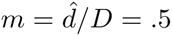. That is, the correlation length parameter estimated by the approximate model is half of the correlation length of the true model.

From these results, the parameter values *k* = 4, *m* = .5 (or any *k* > 2) can be substituted in the expression for *â,* to obtain:

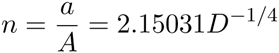

That is, the amplitude parameter of the approximate model is dependent on both the true amplitude parameter as well as the correlation length. The relationship is shown in Figure 3. This parameter could be *n* = .38 − 1.44 scaled from the true amplitude parameter for *D* = 5 − 1000*nm*, relevant scales for membrane protein clusters.

Now, the average number of points per cluster:

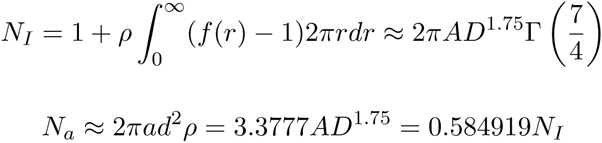

That is, the approximate model underestimates the average number of points per cluster by over 40%.

**Figure 2:**
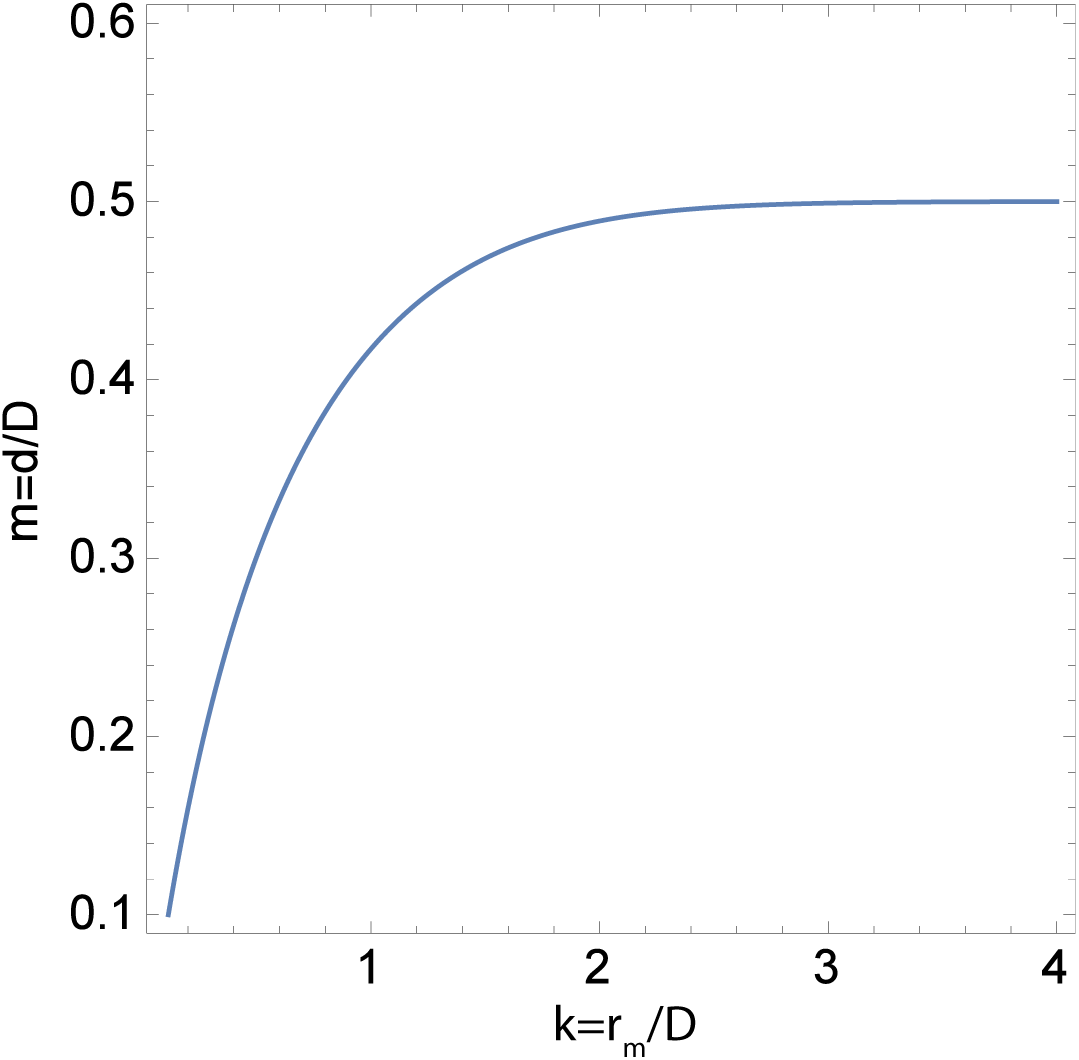
Contour plot of *k* = *r_m_*/*D* vs *m* = *d*/*D* for Ising model. *r_m_* is the distance value to which the Least Squares sum is taken. After ≈ *r_m_* > 2*D*, the *m* value is fixed at .5.

**Figure 3:**
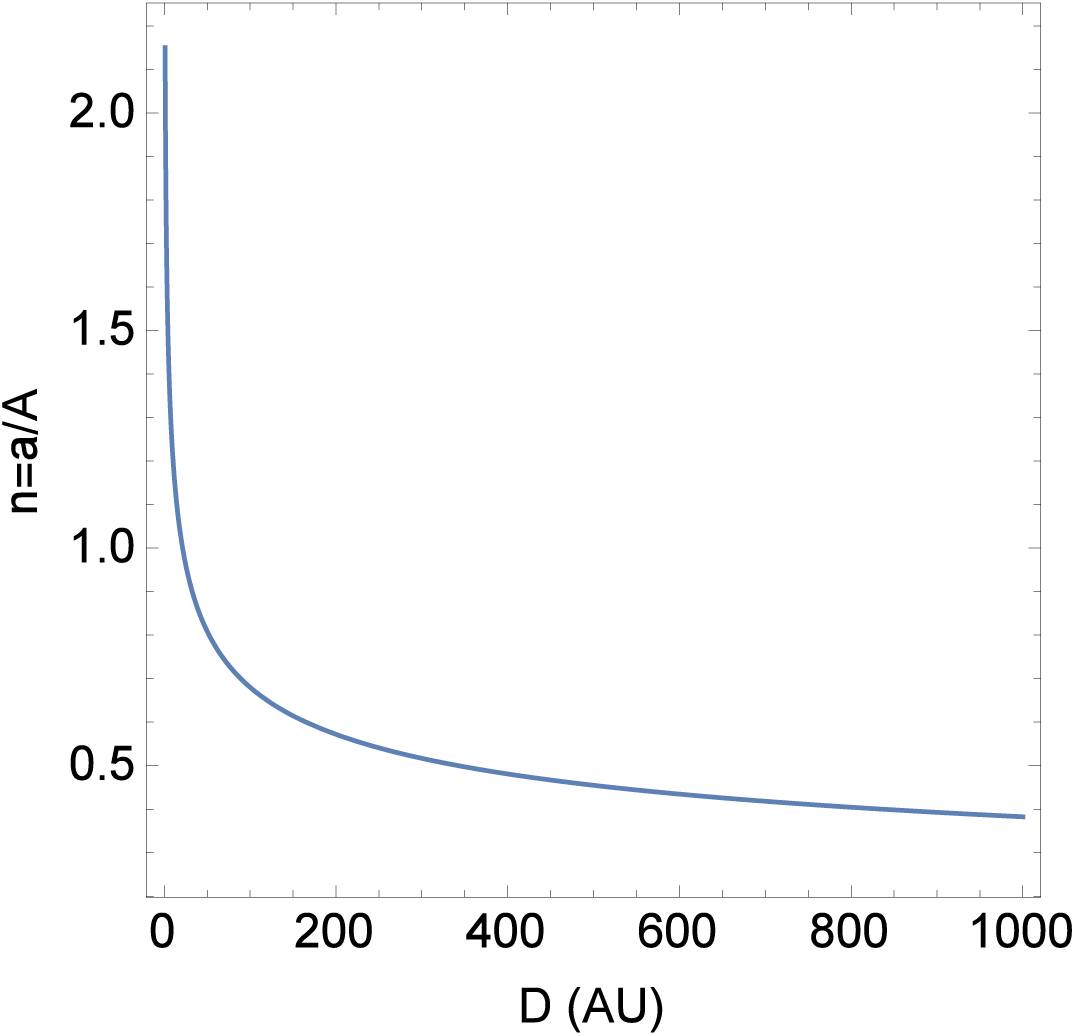
Plot of *D* vs *n* = *a*/*A*, at *k* = 4, *m* = :5.

**Figure 4:**
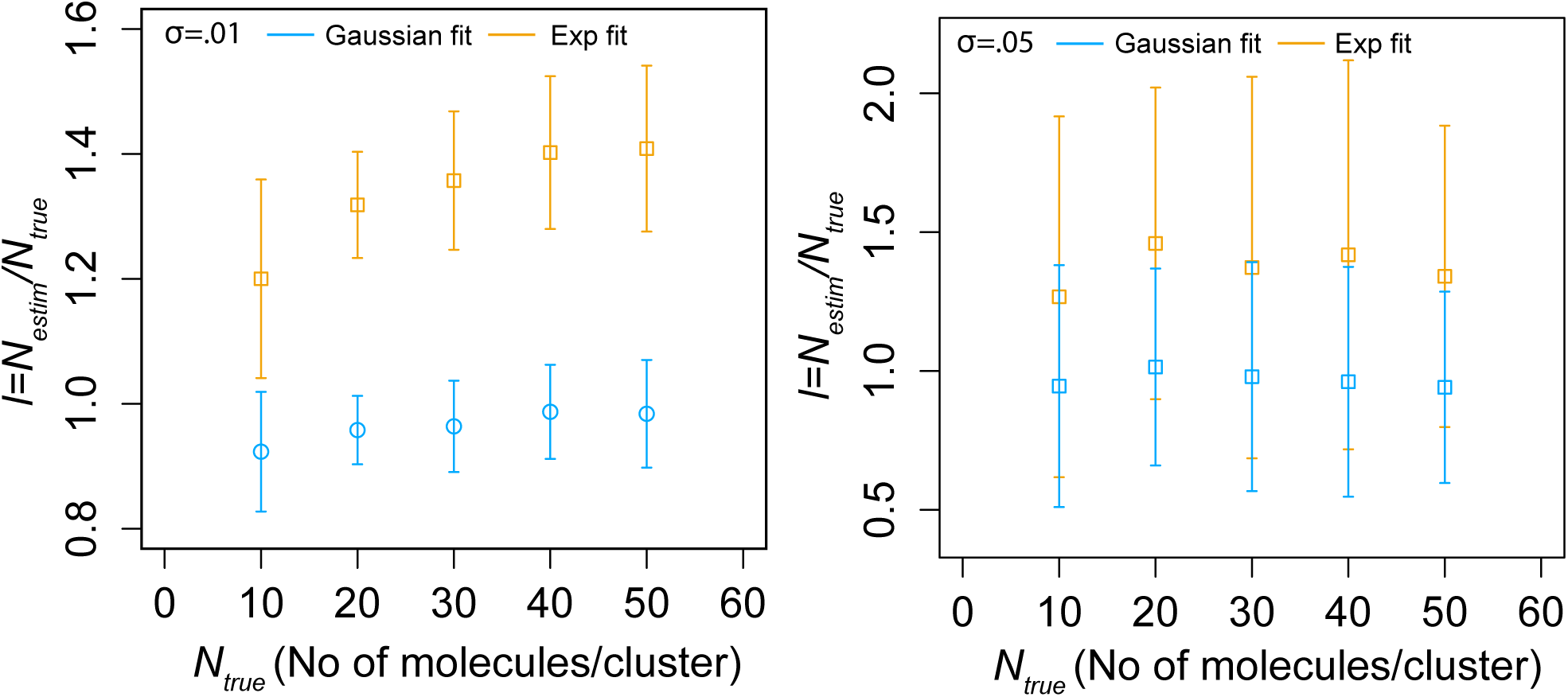
Comparison of fitting empirical PCF of Gaussian clusters to (1) exponential PCF *g_a_* and (2) theoretical PCF of Gaussian clusters, for different true cluster *σ*. Figure 5b in Main Text shown with error bars(*σ*).

### 5 Case of power law PCF

In the case of the PCF 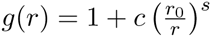, assuming *s* ≠ 1,

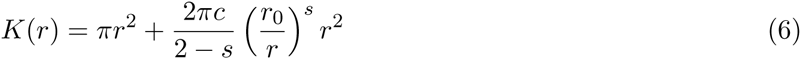

for *s* < 2.

*A* in (10) of Main Text will be 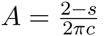. Using (10), we get:

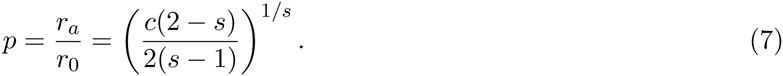

A plot of this equation for different *s* is shown in Figure 5. It can be seen that *p* varies across orders of magnitude based on values of *s* and *c*.

**Figure 5:**
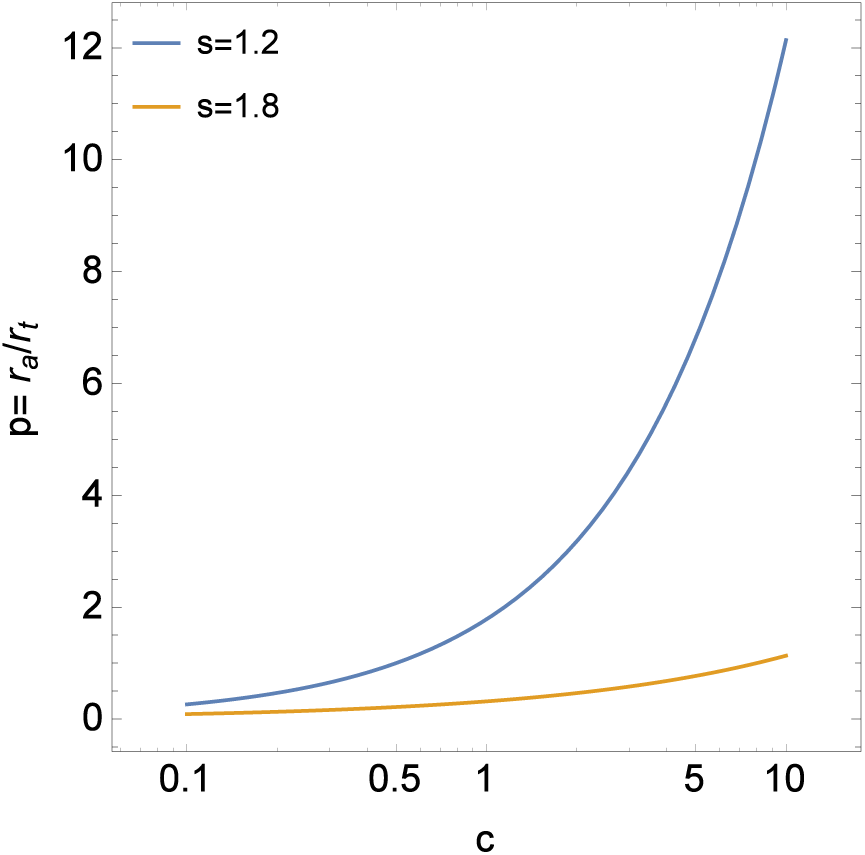
Ratio of radius of maximal aggregation to true cluster size parameter 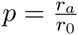 for power law PCF.

